# Molecular characterization of breast and lung tumors by integration of multiple data types with sparse-factor analysis

**DOI:** 10.1101/183582

**Authors:** Tycho Bismeijer, Sander Canisius, Lodewyk Wessels

## Abstract

Effective cancer treatment is crucially dependent on the identification of the biological processes that drive a tumor. However, multiple processes may be active simultaneously in a tumor. Clustering is inherently unsuitable to this task as it assigns a tumor to a single cluster. In addition, the wide availability of multiple data types per tumor provides the opportunity to profile the processes driving a tumor more comprehensively.

Here we introduce Functional Sparse-Factor Analysis (funcSFA) to address these challenges. FuncSFA integrates multiple data types to define a lower dimensional space capturing the relevant variation. A tailor-made module associates biological processes with these factors. FuncSFA is inspired by iCluster, which we improve in several key aspects. First, we increase the convergence efficiency significantly, allowing the analysis of multiple molecular datasets that have not been pre-matched to contain only concordant features. Second, FuncSFA does not assign tumors to discrete clusters, but identifies the dominant driver processes active in each tumor. This is achieved by a regression of the factors on the RNA expression data followed by a functional enrichment analysis and manual curation step.

We apply FuncSFA to the TCGA breast and lung datasets. We identify EMT and Immune processes common to both cancer types. In the breast cancer dataset we recover the known intrinsic subtypes and identify additional processes. These include immune infiltration and EMT, and processes driven by copy number gains on the 8q chromosome arm. In lung cancer we recover the major types (adenocarcinoma and squamous cell carcinoma) and processes active in both of these types. These include EMT, two immune processes, and the activity of the NFE2L2 transcription factor.

In summary, FuncSFA is a robust method to perform discovery of key driver processes in a collection of tumors through unsupervised integration of multiple molecular data types and functional annotation.

**Author Summary:** In order to select effective cancer treatment, we need to determine which biological processes are active in a tumor. To this end, tumors have been quantified by high dimensional molecular measurements such as RNA sequencing and DNA copy number profiling. In order to support decision making, these measurements need to be condensed into interpretable summaries. Such summaries can be made interpretable by connecting them to biological processes.

Biological process activity is continuous and multiple biological processes are taking place in a single tumor. Therefore, the biological processes associated with a tumor are misrepresented by clustering, which tries to put every tumor in a single cluster. In the method introduced in this paper (funcSFA), molecular measurements are summarized into a small number factors. A factor is a continuous value per tumor that aims to represent the activity of a biological process.

When applied to breast and lung cancer, funcSFA identifies factors covering well known biology of these tumor types. FuncSFA also finds novel factors covering biology whose importance is not yet widely recognized in these tumor types. Some of the factors suggest treatment opportunities that can be further investigated in cell lines and mice.

## Introduction

Cancer is a heterogeneous disease, both at the molecular level and in response to treatment. If we can better understand the variation between tumors, we may get a better understanding of why tumors respond differently to treatment. This could, in turn, lead to better treatment selection for patients.

To chart the variation across tumors, projects such as The Cancer Genome Atlas (TCGA) have collected a variety of molecular data from thousands of tumors [1–3]. Analyses of these data provide a better understanding of the underlying biological processes associated with the cancer. For example, recurrent copy number aberrations or recurrent point mutations may reveal the drivers of carcinogenesis. Complementary to this, RNA expression or protein phosphorylation can reveal downstream changes involving many genes, even if the upstream driver of those changes is unknown. Hence, the different data types are reflections of the same biological state, yet each of them encodes information not present or only partially present in the others. Therefore, a comprehensive characterization of the molecular variation across tumors requires the integration of multiple data types.

A popular approach to characterizing tumors is clustering of RNA expression data. Examples include the PAM50 subtypes [4] in breast cancer and the consensus subtypes in colorectal cancer [5]. Since these subtypes are only based on the RNA expression of tumors, they will fail to capture differences in tumors that are more clearly, or even exclusively observed on other molecular levels, such as protein expression, or DNA copy number.

Integrative clustering approaches such as Bayesian consensus clustering [6], patient specific data fusion [7] and iCluster [8] do take multiple data types into account. However, clusters are unsuitable models of biological processes for at least two reasons. First, a biological process can be activated in multiple contexts and multiple independent biological processes can be active simultaneously. However, as clusters cannot overlap, it becomes a challenge to represent this variation in a discrete clustering. For example, immune infiltration occurs in both ER+ and ER− negative breast tumors, but once a tumor is assigned to the ER+ cluster it cannot be assigned to an immune cluster that spans all breast cancer tumors. Second, the variation in activity of a biological process is often more complex than can be captured by a simple distinction between absent or present. Instead, it is more naturally expressed along a continuous scale of activity levels. This cannot be captured by discrete clusters.

Paradigm [9] improves upon the abovementioned approaches by integrating multiple data types to infer activity levels of biological processes. Activity levels of biological processes in tumors are assigned independently of each other, avoiding the limitation of cluster analysis. To estimate these activity levels, Paradigm leverages existing knowledge available from pathway databases. A limitation of this approach is that using existing knowledge a priori limits discovery of new biological processes. More importantly, it also limits the discovery of biological processes in new contexts (e.g. tumor types) because activity of a process in a new context might involve a set of genes that is only partially overlapping with the genes currently annotated to that process.

Here we introduce FuncSFA, a sparse-factor analysis with a tailored gene-set enrichment analysis (GSEA) [10] that integrates multiple data types to provide both a continuous characterization and a functional interpretation of the variation across tumors at the molecular level (Fig 1). The sparse-factor analysis identifies factors explaining variation in multiple data types such as RNA expression, protein expression, and DNA copy number data. Subsequently, the factors are interpreted and linked to known biology using a gene-set enrichment analysis of the factors on the RNA expression data. The interpretation obtained from the gene-set enrichment analysis is validated by comparison of the genes, epitopes and copy number aberrations in the factor to external resources. Together this not only provides insight into variation across tumors but also into the biology underlying the molecular data.

The sparse-factor analysis is based on a reinterpretation of the mathematical framework behind the iCluster method [8, 11]. Our reinterpretation improves upon the iCluster method in several key aspects. First, through proper factor rescaling and more efficient optimization approaches, we ensure convergence of FuncSFA on multiple molecular datasets that have not been pre-matched. An example of pre-matching is in the iC10 subtypes [12], where the authors only selected genes where RNA expression correlates with DNA copy number. Second, in contrast to iCluster, FuncSFA does not assign tumors to discrete clusters, but identifies the dominant driver processes across multiple molecular data types and across all tumors, and then represents, for each tumor, the spectrum of processes active in that tumor. Taken together, FuncSFA represents, for the first time, a robust method to perform discovery of the key biological processes driving the most important phenotypic differences across a set of tumors through unsupervised integration of multiple molecular data types.

We applied FuncSFA to the TCGA breast and lung cohorts and identified 10 factors in each tumor type. As breast cancer is very well characterized, it served as a positive control in the sense that we could fully identify the known intrinsic subtypes in breast cancer. Uniquely, this characterization integrates the intrinsic subtypes and the activity of Epithelial to Mesenchymal Transition (EMT) and Immune processes, each represented by an independent factor. In lung cancer, which has remained largely uncharacterized, we also identified an EMT and Immune factor, as well as a factor associated with the main lung subtypes—Adenocarcinoma and Squamous Cell Carcinoma. We also identified a factor which captures the activation of the transcription factor NFE2L2. Here the power of integration of multiple data types is highlighted by the fact that the activity of this factor is associated with mutations in NFE2L2 as well as its inhibitor KEAP. We expect that the identified factors not only provide a more complete characterization of the biological processes active in the different tumors, but will also provide a starting point for the development of better treatment strategies.

**Fig 1.**
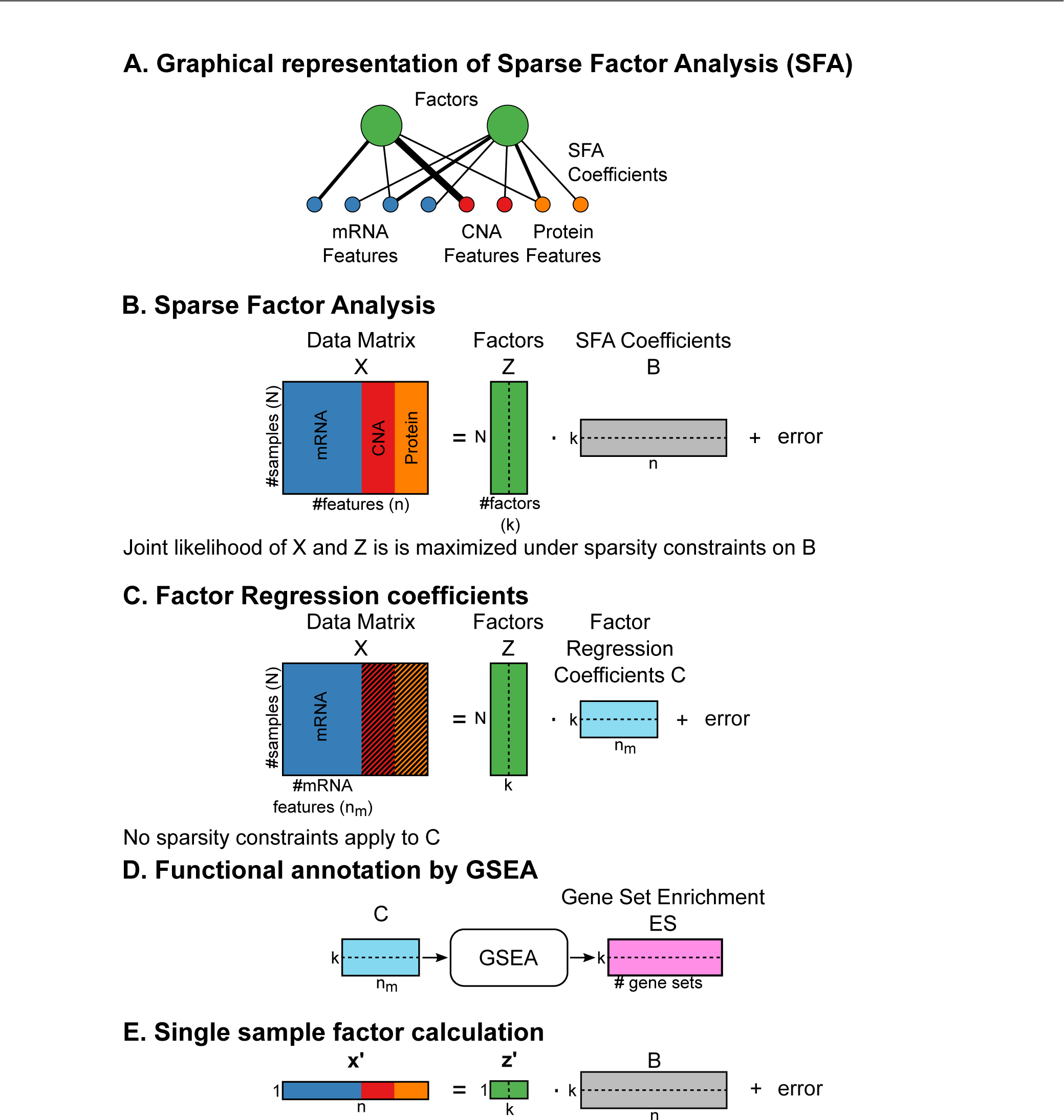
Overview of FuncSFA. A: Graphical representation of Functional Sparse-Factor Analysis (FuncSFA). The green circles represent the factors, and the red, blue and yellow circles at the bottom represent the observed variables, with the colors representing the data types and each circle representing an individual variable (i.e. the expression of a gene or protein, or the copy number of a gene). The black lines connecting the individual variables to the factors represent the regression coefficients. B: Graphical representation of the mathematical concepts of SFA with X representing the N × n data matrix, Z the N × k obtained factor matrix and B the k × n factor coefficients. C: Graphical representation of the computations of the factor expression coefficients. The coefficients represented by the k × n_m_ matrix C are obtained by regressing the N × n_m_ RNA expression matrix, X_m_, on the N × k factor matrix Z. D: The gene-set enrichment analysis designed to assign biological processes or pathways to the obtained factors. E: Application of the factors to determine the activity of the factors (or associated biological processes) in a new tumor. (N: number of tumors; n: number of features; k: number of factors; n_m_: number of mRNA features; Z: factor matrix; X: data matrix (concatenation of mRNA, copy number and Reverse Phase Protein Array (RPPA) data); B: Sparse factor coefficients; C: Factor regression coefficients; GSEA: Gene-set enrichment analysis).

## Results

### FuncSFA

FuncSFA consists of three components. The first component performs the sparse-factor analysis to obtain the factors. In the second component, Gene Set Enrichment Analysis is employed to interpret the obtained factors in terms of the possible biological processes they represent. The last component allows the application of the obtained factorization to a new sample in order to reveal the biological processes likely giving rise to the molecular profiles observed for that sample. In this section we discuss each of these components.

### Sparse-factor analysis

In our sparse-factor analysis, we assume that each tumor type is characterized by a set of key driving factors, or biological processes, that give rise to molecular phenotypes. These driving factors cannot be measured directly, but are observed indirectly through noisy measurements of multiple molecular data types such as mRNA expression, copy number aberrations and protein expression and modification. The challenge is to employ all these measured data types simultaneously to identify this unobserved structure in the data.

Sparse-factor analysis addresses this challenge by modeling the unobserved structure by a relatively small number of continuous factors— the green circles in Fig 1A. These factors, in turn, explain the observed molecular data in a linear regression that links each molecular data type to the factors. The red, blue and yellow circles at the bottom of Fig 1A represent the observed variables, with the colors representing the data types and each circle representing an individual variable. For example, in ERBB2 (also known as HER2) positive breast cancer, the ERBB2 factor would represent the activation of the ERBB2 pathway while the measured copy number, protein and mRNA expression changes will be modeled as appropriately chosen regression functions of the ERBB2 factor.

The factors are identified by maximizing the joint likelihood of the the measured data and the factors, selecting factors and regression coefficients explaining the measured data. This is accomplished in a methodology that is akin to probabilistic principal component analysis (PCA) [13]. We employ elastic net regularization [14] to create parsimonious models by enforcing sparsity on the regression coefficients associated with the factors, as introduced by iCluster [8]. In Fig 1A the regression coefficients are represented by the black lines connecting the individual variables to the factors, while Fig 1B depicts the mathematical operations associated with the sparse-factor analysis. Consequently, a factor is defined by the subset of molecular variables that contribute to that factor as well as the regression coefficients that model the degree to which each molecular variable contributes to the factor.

Sparsity of regression coefficients induced by the elastic net penalty improves data integration by preventing one data type (especially mRNA expression) from dominating the analysis. A larger penalty can be applied to mRNA expression to the other data types, keeping mRNA expression under control. Sparse weights are also biologically plausible as a biological process might involve the altered expression of thousands of genes but is unlikely to be driven by all genes.

The joint likelihood is maximized using an expectation maximization algorithm. This algorithm improves over the iCluster2 algorithm [11] by rescaling the factors to unit variance and by estimating coefficients with coordinate descent [15]). On the TCGA breast cancer dataset our algorithm converged faster and to better solutions than the iCluster2 algorithm (S1 Fig).

A very important feature of FuncSFA is that, unlike iCluster, we do not force a tumor assignment to discrete clusters, but represent the level of activity of each of the driving factors in a given tumor. Consequently, each tumor can be a 'member’ of multiple factors, and this 'membership’ can also vary in strength.

### Gene-set enrichment analysis

While sparse-factor analysis efficiently identifies the hidden driving factors, it remains challenging to directly attach biological interpretation to the identified factors. In some cases, such as ERBB2 pathway activation, this may be straightforward, but in many other cases this remains very challenging. One of our important contributions is the development of a gene-set enrichment analysis tailored to the results of sparse-factor analysis.

The first step of the gene-set enrichment analysis is to regress the factors on the gene expression matrix (Fig 1C). This may sound counter-intuitive, as the purpose of the SFA is to pinpoint the genes contributing to the factor, hence revealing the underlying drivers. However, the motivation for this is two-fold. First, the sparsity constraints on the coefficients introduce zeros in the coefficients. Although this is beneficial for human interpretation and driver identification, genes with zero coefficients can not be ranked in terms of their contribution to the factors. This is a major complicating factor as such a gene ranking is an essential part of the gene-set enrichment analysis. Second, RNA expression remains a data type that captures most of the variation in the cell, and can, as such be quite informative regarding the activity of biological processes and hence for the interpretation of the factors. So, having gone through the process of identifying driving factors that are robust, in the sense that they are common to all data types, we employ the regression of the factors to the complete RNA expression data set to identify all genes with expression patterns that show association with the factors. This association is captured in the regression coefficients (Matrix C in Fig 1C). We then normalize each coefficient by the gene standard deviance to obtain the ‘factor expression coefficients’.

The second step of the gene-set enrichment analysis is to rank the genes based on the factor expression coefficients and compute the enrichment statistic for every gene-set factor pair (Fig 1D) [10]. We determined statistical significance of the enrichments by a sample permutation test [10]. The enrichment results provide input for a manual curation and verification process to identify the most likely biological process that gives rise to the identified factors.

### Single-sample analysis

Finally, in order to determine the activity of the identified factors in a new unseen tumor, one simply solves the same equation with the new tumor(s) while keeping the sparse factor coefficients fixed, as depicted in Fig 1E. Importantly, this can be done with the gene expression data only, allowing easy translation of the factors to other datasets.

### Application to breast and lung cancer

We have applied FuncSFA to the breast cancer [1] and lung cancer [2, 3] data sets from TCGA. Breast cancer is arguably the most exhaustively subtyped type of cancer, and hence serves as a very good positive control for FuncSFA [1, 12, 16, 17]. Lung cancer has not been studied so extensively, even though at the moment it is the cancer type resulting in the highest number of deaths per year. For this reason, there is a large unmet need for the discovery of new biomarkers which could open up new treatment modalities. From a technical perspective, TCGA includes large numbers of tumors for both cancer types, which ensures that the parameter estimates produced by FuncSFA are reliable.

For both datasets, we used three data types: DNA copy number, protein expression measured by RPPA, and RNA expression. First, we included DNA copy number at 162 (breast) or 213 (lung) frequently aberrated loci from SNP6 arrays as identified by RUBIC [18]. This DNA copy number data set is clearly important as it captures many copy number events that may have a functional role in oncogenesis. Second, we included protein expression and modification recorded by RPPA with 195 (breast) or 216 (lung) protein epitopes. These measurements capture the activity of key signaling events in pathways which play a central role in many cancer types, including those under study here. Third, we included RNA expression of the 1000 most variable genes as measured by RNAseq. We selected the 1000 most variable genes to reduce noise in the data and to reduce the complexity of the model. We included RNA expression as it is arguably the most comprehensive data type, capturing the downstream effects of many upstream events such as signaling pathway activity or activation of cancer genes by, for example, genetic events. Although FuncSFA allows the inclusion of many data types, we have not included mutation data. The binary nature of this data does not directly fit the factor analysis model we employed (see Discussion and Methods).

For both breast and lung cancer, we employed the abovementioned data types and applied FuncSFA. We specifically set out to find the ten strongest factors. We chose this number of factors based on the following argumentation. First, when increasing the number of factors up to at least twenty, one can discover more detailed processes showing activity, provided that a sufficiently large sample size is employed. However, as the number of factors increases, the strongest factors remain the same (S2 Fig). Second, most subtyping approaches that have been applied to date revealed ten or fewer subtypes: five intrinsic subtypes in breast cancer [19, 20] ten IC10 subtypes in breast cancer [12], four consensus subtypes in colorectal cancer [5], four in squamous cell carcinoma [21] and three in lung adenocarcinoma [22]). Therefore, our assumption was that ten factors would be sufficient to capture the strongest (known) factors, while leaving room for the discovery of new biological processes without running the risk of compromising the robustness of the factors being discovered by having a too large factor-to-sample-size ratio.

### Breast Cancer

We applied FuncSFA to the breast cancer data employing 10 factors based on the arguments given above. We performed functional annotation of the factors, as outlined above and in the methods section, employing, amongst others, the coefficients obtained from the sparse-factor analysis, as well as the gene-set enrichment for pathways and biological processes depicted in the supplement (S4 Fig and S1 Table). This resulted in the following 10 factors: ER (Estrogen Receptor), Normal-like, Basal, HER2, Luminal Proliferative, 8q-gained, Technical-RNA, Technical-RPPA, EMT (Epithelial to Mesenchymal Transition) and Immune. The strongest sparse-factor analysis coefficients (from the B matrix in Fig 1B) for the three data types are represented in Fig 2. We will first provide some general observations of the results and then provide a detailed description and analysis of each factor.

From the coefficients depicted in Fig 2, we make the following global observations. First, the ER factor is strongly associated with both mRNA and protein expression of ESR1, GATA3, PGR and AR, as expected. Second, the EMT factor shows strong association with THBS2 and COL11A1 expression—the genes identified by Anastassiou and colleagues [23] (see also the more detailed description of the EMT factor below). In addition, the EMT factor shows association with many collagens. Third, the HER2 factor shows the expected strong association with ERBB2 copy number gain and protein upregulation, GRB7 copy number gain as well as EGFR protein upregulation. Fourth, the Basal factor shows strong association with the RNA expression of basal keratins and finally, 8q-gained shows concordant copy number and expression changes for a number of genes located on Chromosome 8q, including SGK3, MYC and TP52INP1.

Fig 3A depicts, per factor, for all ten factors, the amount of explained variation for the three data types: gene expression, copy number and RPPA. Most factors explain variation in each of the data types, with the largest proportion of the variation explained by a given factor mostly being the variation in gene expression. There are a few exceptions. The HER2, Luminal proliferative and 8q-gained factors, explain more variation in copy number data than in RNA expression data. For HER2 and 8q-gained tumors this is not surprising as they are clearly copy number driven (Fig 2 and 4). Most strikingly, the Techical-RPPA factor explains a very large proportion of the variation in the RPPA data. However, as the name indicates this factor most likely captures technical variation in the RPPA data, as this variation is not reflected in the other data types, and as the gene-set enrichment analysis does not reveal a clear functional enrichment associated with this factor. Similarly, the Technical-RNA factor explains technical variation in the RNAseq data. The ER factor is unique in the sense that it is the single factor that explains most variation in RNA expression and RPPA data (apart from the Technical-RPPA factor). This is not unexpected as ER signaling arguably drives the most important subtype distinction in breast cancer: the one between ER-positive and ER-negative tumors. In the remainder of this section, we will discuss each of the identified factors in greater detail.

**Fig 2.**
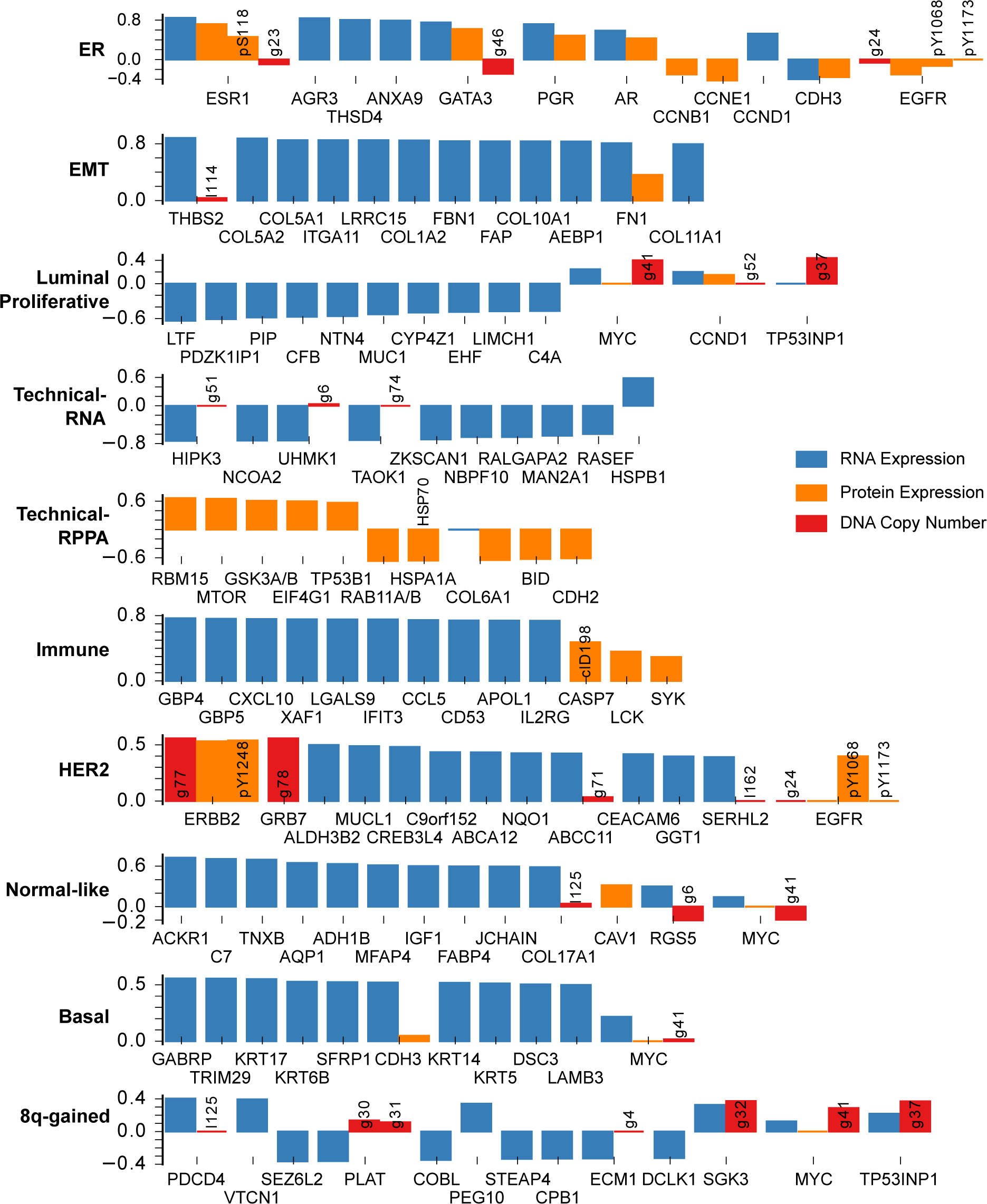
The strongest sparse-factor analysis coefficients for the breast cancer data set for each of the three data types and all ten factors. The height of the bars shows the values of the coefficients. If a gene is strongly associated with a factor, we show all coefficients of that gene in the model. RNA expression coefficients are shown in blue. Protein expression coefficients are shown in orange. Any modifications of an epitope are noted in a short text description: pX = phosphorylated at residue X; clX = cleaved at residue X. DNA copy number coefficients are shown in red. Numbers refer to the recurrently aberrated loci in S3 Table. Recurrent gains are prefixed with a g, losses with an l.

### Intrinsic subtypes are covered by factors

The intrinsic subtypes proposed by Perou and Sørlie represent one of the earliest and most widely used subtypings of breast cancer [17, 19, 20]. This discrete classification of breast cancer in five subtypes (Her2-enriched, Basal-like, Luminal A, Luminal B and Normal-like) is typically performed by applying a nearest centroid classifier to the RNA expression profile of the tumor to be subtyped. To perform this classification based on mRNA expression, Parker and colleagues developed a a 50 gene signature, the so-called PAM50 [4]. As the PAM50 subtyping is widely acknowledged as a gold standard in breast cancer subtyping, we set out to check whether the variation captured by the PAM50 subtyping is also recapitulated by the FuncSFA factors. To this end we applied the PAM50 subtyping to the breast cancer cohort and compared the PAM50 subtyping to the variation captured by the FuncSFA factors. Fig 3B.1 depicts a t-SNE map of the breast cancer tumors with the PAM50 subtype assignment indicated by the colors. Four FuncSFA factors capture the variation in the PAM50 subtypes.

First, the ER factor is associated with both mRNA and protein expression of ESR1, AR and PR (Fig 2), and is enriched for gene signatures of ESR1 expression and the luminal subtype of breast cancer, which, in turn, is characterized by ESR1 overexpression (S4 Fig and S1 Table). This suggests that this factor represents the continuous variation in ESR1 expression in breast cancer, which is typically dichotomized to define ER+ and ER− tumors. This is confirmed by comparing the PAM50 subtypes with the ER factor. Indeed, the ER factor is strongly associated with the ER+ (Luminal A, Luminal B and Normal-like) and ER− (Basal-like, HER2-enriched) subtypes. Specifically, classification into the ER+ and ER− classes based on the ER factor results in an AUC of 0.98. The strong association between the ER factor and the ER+/ER− PAM50 subtyping is also strikingly visible in Fig 3B.2, where the same tumor positions as in Fig 3B.1 are maintained whilst the tumors are colored according to the value of the ER factor.

Second, the HER2 factor shows large SFA coefficients for ERBB2 protein expression and copy number gain (Fig 2). The gene-set enrichment analysis shows enrichment for signatures of ERBB2 amplification and the HER2-enriched PAM50 subtype. Importantly, this factor is strongly associated with the amplicon on Chromosome 17 harboring ERBB2 and GRB7, as evidenced by the large coefficients identified for this amplicon in the factor analysis (Fig 2 and 4). Taken together, this suggest that the HER2-factor indeed identifies the HER2+ tumors. This is confirmed by the strong association of this factor with the HER2-enriched PAM50 subtype (Fig 3B.3, AUC=0.96) and by the large weights of the HER2/GRB7 locus on DNA copy number (Fig 4).

Third, the Luminal-Proliferative factor shows enrichment for signatures representing the cell cycle, proliferation and high-grade tumors in the gene-set enrichment analysis. This factor shows strong association with the PAM50 Luminal A and Luminal B subtypes (AUC of 0.78), with the factor being predominantly low in Luminal A and high in Luminal B (Fig 3B.4). Notice that, in contrast to the ER and HER2 factors which are continuous but show bimodality, this factor is more unimodal (S3 Fig). In addition, the copy number weights associated with this factor shown in Fig 4, show the 8q and 17q gains and 8p losses characteristic of some Luminal B tumors [12].

**Fig 3.**
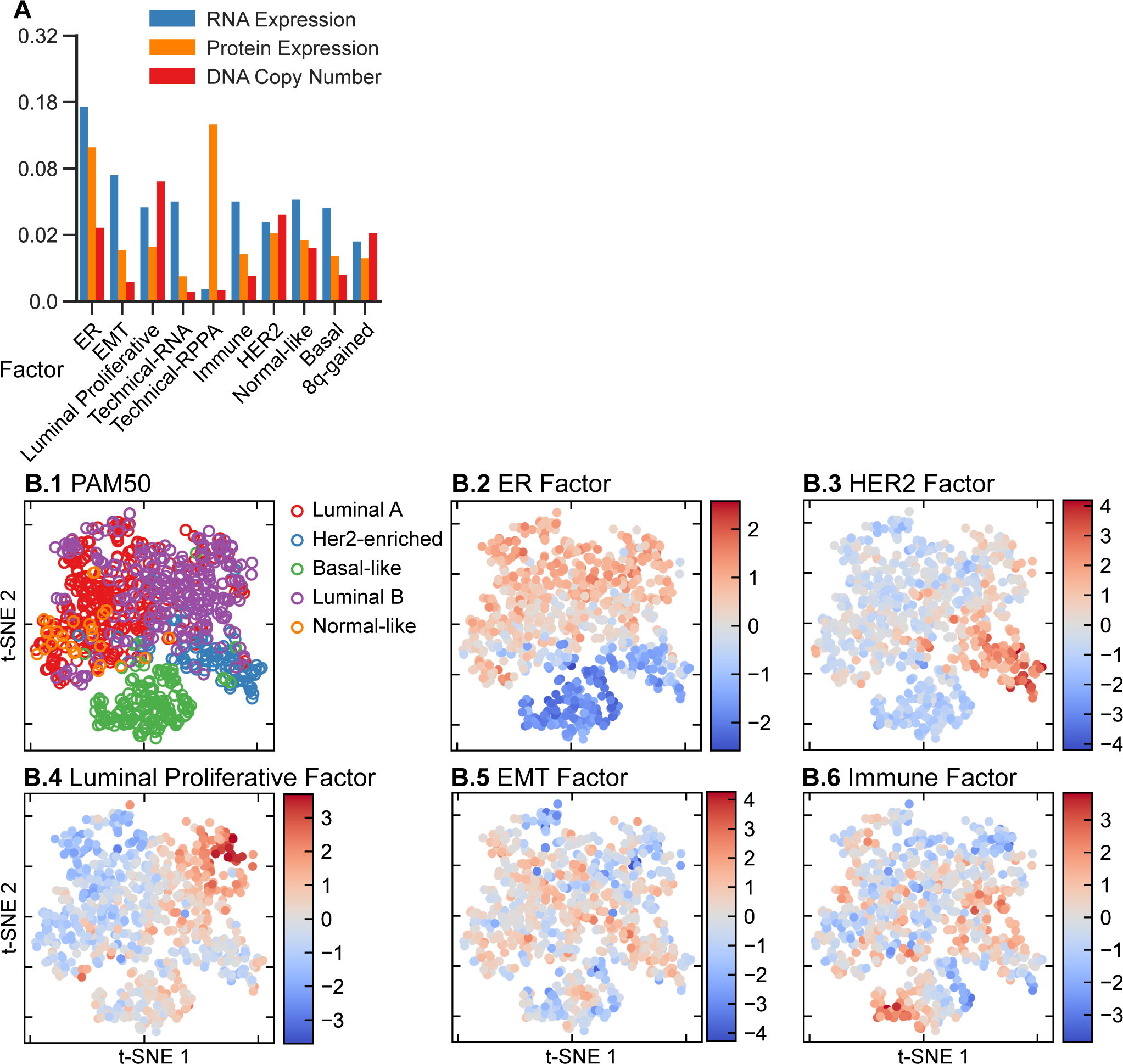
Sparse-factor analysis on the TCGA breast cancer data set. A: Explained variation per data type and factor. B: The top-left panel shows a t-SNE map of the tumors with the different colors showing PAM50 subtypes. The remaining panels show the tumors in the same positions as the PAM50 map, but colored according to the value of the represented factor in each tumor.

**Fig 4.**
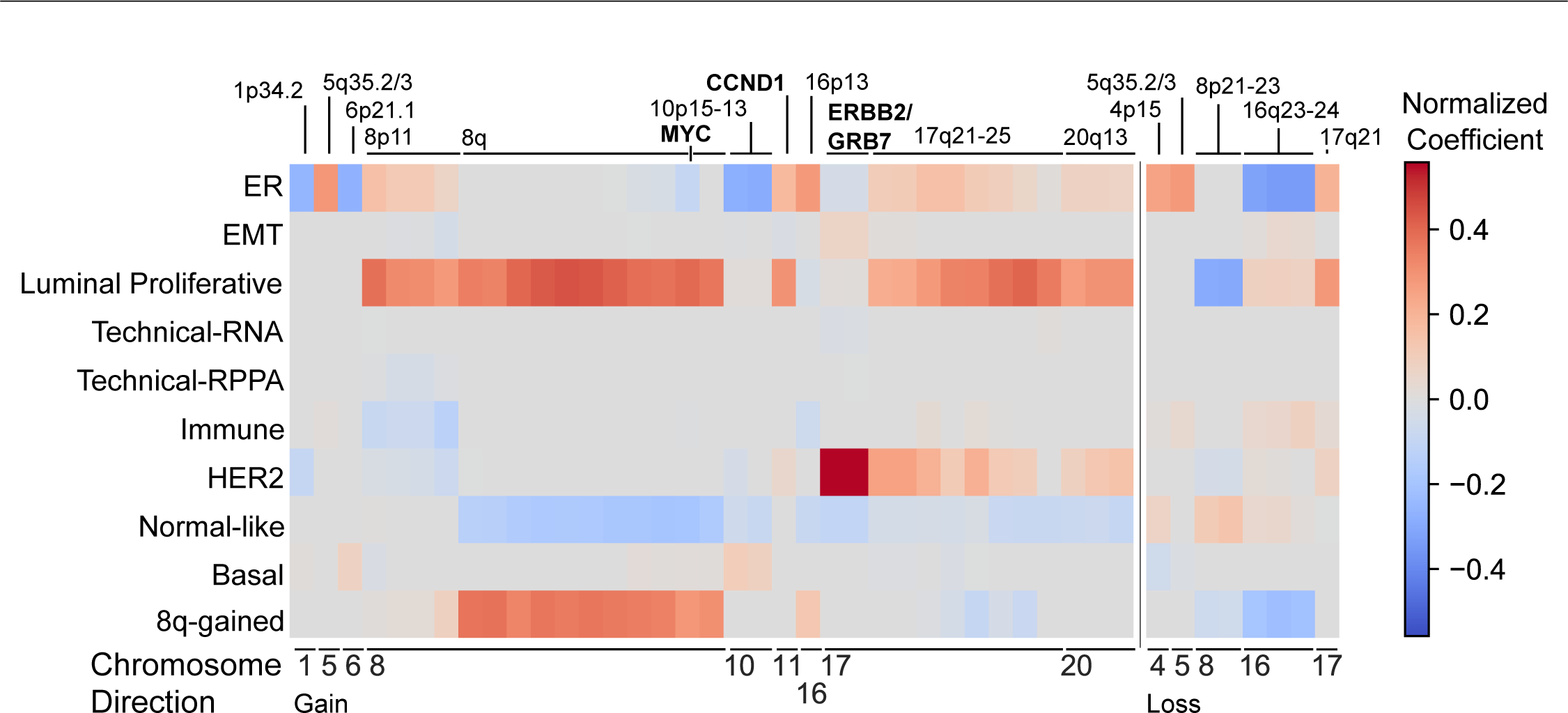
Copy number and factors in the TCGA breast cancer dataset. Normalized coefficients representing the contribution of DNA copy number aberrations to the factors. Specifically, the coefficients represent the contribution of recurrently gained (left) or lost (right) copy number regions identified by RUBIC to the factors represented in the rows. Recurrently aberrated copy number regions are annotated with chromosomal bands or putative driver genes in the region.

Fourth, gene-set enrichment analysis and ROC-curves show that the Normal-like factor is associated with PAM50 normal-like tumors (AUC=0.92) and tumors with a lobular pathology (AUC=0.84). Finally, the Basal factor is associated with expression of the basal keratins KRT5, KRT6B, KRT14 and KRT17 (Fig 2) and appears only within the triple-negative tumors (S6 Fig). The Basal factor is very different from the Basal-like intrinsic subtype. The latter encompasses almost all triple negative tumors and it is not directly related to basal cells, a relationship we have shown for the Basal factor based on the expression of basal keratins. Taken together, we can conclude that the ER, HER2, Luminal Proliferative and Normal-like factors collectively capture most of the variation in the well-known breast cancer intrinsic subtypes.

### EMT factor

EMT is a process frequently associated with cancer [24] and it involves multiple regulators, including SNAI1, SNAI2, TWIST1 and ZEB1 as well as their targets [25]. The gene-set enrichment analysis revealed an association of one of the factors with EMT and the extracellular matrix (S1 Table and S4 Fig). Of the known regulators of EMT, only SNAI1 is on the RPPA array, and its protein expression is not associated with this factor. Gene expression of the EMT regulators is correlated to the factor (Spearman correlation, SNAI1: ρ=0.22, SNAI2: ρ=0.63, TWIST1: ρ=0.42, ZEB1: ρ=0.69). More generally, quite a number of gene expression signatures have been developed to detect various forms of EMT (For example, see [26]). Based on the gene-set enrichment analysis, the strongest association of the EMT factor is with a consensus EMT signature proposed by Anastassiou and colleagues [23]. They compiled a pan-cancer EMT signature from multiple public data sets by comparing metastatic with non-metastatic tumors [27]. As an EMT-like expression profile can be associated with stromal contamination in a tumor sample, Anastassiou and colleagues profiled human tumors from a PDX model on a microarray with species-specific probes. This enabled the removal of the mouse stromal signal and revealed that the signature is tumor specific. A strong correlation between the EMT factor and the sum of the two most important genes in this signature (Fig 5A, ρ=0.89) confirms the interpretation of this factor as capturing the specific type of EMT modeled by the Anastassiou signature.

While the claudin-low and metaplastic subtypes of breast cancer have been associated with EMT [28], larger studies failed to confirm the presence of the claudin-low subtype [1]. The t-SNE map of Fig 3B.5 and the boxplots in the supplement (S9 Fig) reveal that this factor is equally strongly represented in all intrinsic subtypes. It is therefore impossible to have an EMT-high subtype that includes all EMT high tumors in a clustering that also captures the intrinsic subtypes, which possibly explains the lack of reproducibility of the claudin-low and metaplastic subtypes. Possibly, the EMT factor captures part of the biology captured by the claudin-low and metaplastic subtypes, while taking the context of other subtypes into account.

**Fig 5.**
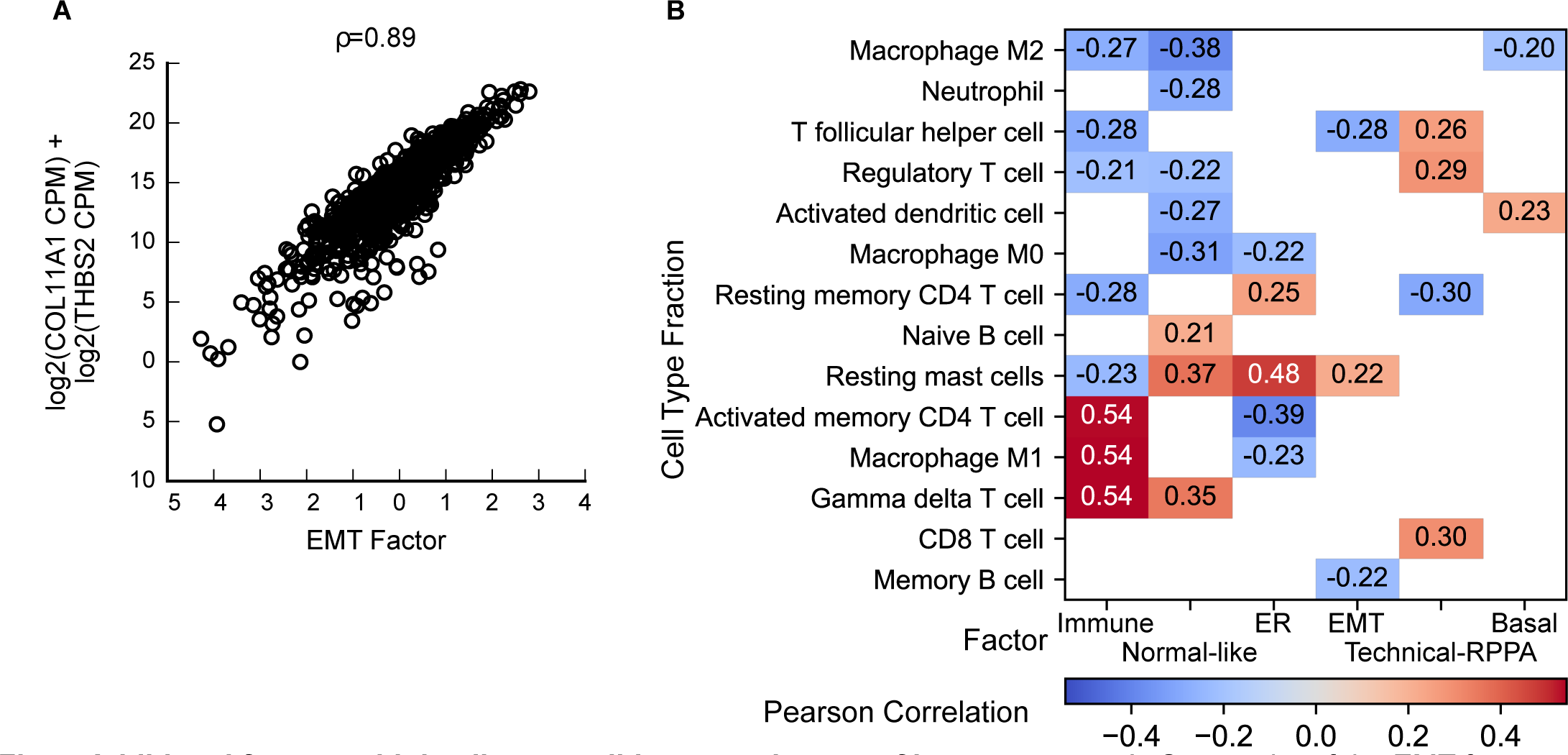
Additional factors add detail over well-known subtypes of breast cancer. A: Scatterplot of the EMT factor versus the sum of the RNA expression of COL11A1 and THBS2 (CPM: count per million). B: Pearson correlation (ρ) between the factors and cell type fractions. Only significant correlations (p<0.05, |ρ|>0.2) are shown.

### Immune factor

The Immune factor shows enrichment for Interferon-Alpha Response and other immune related signatures (S1 Table and S1 Table). In order to shed further light on this factor we employed publicly available Cibersort scores [29] to estimate the immune cell fractions in all the breast tumors, based on their RNA expression profiles [30]. Briefly, Cibersort employs RNA expression profiles of 22 pure immune cell types to perform an in silico decomposition of the RNA expression profile of a tumor. The resulting output provides an estimate of the relative abundance of the different immune cell types in the tumor being decomposed. In other words, for every immune cell type, we obtain a profile across all the breast cancer tumors representing the estimated fraction of that cell type in each tumor. We then computed the Pearson correlation (ρ) of these immune cell type profiles with each of the factors. Fig 5B depicts the significant correlations, i.e. only factor and immune cell profile correlations where p<0.05 and |ρ|>0.2. The Technical RPPA, ER, Normal-Like and Immune factor all show more than one significant correlation with an immune cell type. Whereas the ER factor (resting memory DC4 T cells, resting mast cells) and the normal-like factor (γδ T-cells, resting mast cells, naive B cells) both display a more indolent immune profile, the Immune factor clearly shows high positive correlation with active immune cell types and negative correlation with resting immune cells. While the immune factor is quite uniformly present in all breast cancer tumors, there is an enrichment for very large values in the Basal-like tumors which corresponds with previous reports of CD274 (PD-L1) expression patterns [31], and Cibersort inferred immune infiltration patters [32].

Taken together, we have identified 10 factors in breast cancer. A number of these factors (ER, Normal-like, HER2, Basal and Luminal-Proliferative) capture the subtype variation previously described in the Intrinsic subtypes. This serves as a positive control of FuncSFA as it illustrates that it can capture known biological variation. More importantly, FuncSFA also captured additional variation not represented in the Intrinsic subtypes. Specifically, we identified the 8q-gained factor which is associated with gains of the q-arm of Chromosome 8 and loss of the Chromosome 16q23-24 region, as is clear from the DNA copy number weights in Fig 4. In addition, we identified EMT and Immune factors that may lead to a better understanding of the processes playing an important role in breast cancer, while also serving as a starting point for the development of new treatment approaches.

### Lung Cancer

Having established the utility of FuncSFA on the breast cancer data, we moved on to assess FuncSFA on a less well-characterized tumor type. To this end we applied FuncSFA to the lung tumor data from the TCGA in order to further assess its usefulness in uncovering new and clinically relevant biology. We merged the TCGA lung adenocarcinoma and lung squamous cell carcinoma datasets to obtain a data set that is comparable in size to the breast cancer set. We employed the same data types (gene expression, RPPA and copy number) as we employed in the breast cancer analysis. As the dataset is of comparable size to the breast cancer dataset, and as Wilkerson and colleagues defined a total of 7 lung cancer subtypes [21, 22] we set out to identify 10 factors. This allows the capturing of known variation with some room for the discovery of novel subtypes.

FuncSFA identified 10 lung cancer factors: Adenocarcinoma, Mitochondria, DNA replication, Immune, Infiltrating B-cells, NFE2L2, EMT, Translation, BSCC (Basal Squamous Cell Carcinoma) and 8p11-gained. S5 Fig and S2 Table list the factors and the processes and pathways that were significantly enriched in each of the factors based on the gene-set enrichment analysis. The strongest SFA coefficients (the B matrix in Fig 1) for the three data types are represented in Fig 6. As before we will first provide some general observations of the results and then provide a detailed description and analysis of each factor.

The coefficients depicted in Fig 6 reveal the following. First, as before, the EMT factor shows strong association with the Anastassiou signature genes (THBS2 and COL11A1) as well as a number of collagens. Second, the Mitochondrial factor has large SFA coefficients for genes encoded on the mitochondrial DNA while the Translation factor shows strong association with genes encoding ribosomal proteins. Third, the BSCC and 8p11-gained factors show large RPPA coefficients, with a concentration of lowly expressed proteins in the PI3K pathway. However, they do differ in terms of their RNA expression coefficients, and we will provide more elaborate descriptions of these factors below. Fourth, the Immune factor is characterized by large RNA expression coefficients of immunoglobulins. Finally, the Adenocarcinoma factor shows association with the RNA expression of a number of keratins, negative coefficients for both copy number and RNA expression of SOX2 and positive association with both RNA expression and copy number of NKX2-1 (also known as TTF1).

Fig 7A depicts, per factor, for all ten factors, the amount of explained variation for the three data types: gene expression, copy number and RPPA. As in breast cancer, most factors explain variation in each of the data types, with the largest proportion of the variation explained by a given factor mostly being the variation in RNA expression with the exceptions being the 8p11-gained and BSCC factors where the largest portion of the variance explained by these factors is the variation in the RPPA data. It is also noteworthy that these two factors explain a significantly larger portion (≈0.32) of the variation in the RPPA data, compared to the factors that explain the largest portion of the RNA expression variation (≈0.19). The 8p11-gained and BSCC factor activity levels are negatively correlated and have RPPA coefficients of similar magnitude and equal sign (S8 Fig). This means that the RPPA values predicted by these two factors together are much smaller than the RPPA values predicted by one of these factors on its own.

In the remainder of this section, we will first discuss the Wilkerson subtypes that have been proposed for lung cancer. Then we will discuss each of the identified factors in greater detail.

### The Wilkerson molecular subtypes of lung cancer

The molecular subtypes of lung adenocarcinoma and lung squamous carcinoma were defined by Wilkerson and colleagues [21, 22], and also found to be present on the larger TCGA datasets [2, 3]. Fig 7B.2 depicts the Wilkerson subtypes on top of the t-SNE plot derived from the SFA factors. Here every tumor is colored according to its subtype membership. Globally, all SFA factors are associated with these subtypes (p < 10^-6^ for every factor, Kruskal-Wallis test). However, in contrast to breast cancer, it was not possible to perform a one-to-one mapping from the SFA factors to the Wilkerson subtypes. This can most likely be explained by the fact that none of the Wilkerson subtypes have (yet) been associated with a driver event, unlike breast cancer where ESR1 and ERBB2 have been identified as strong drivers giving rise to specific subtypes of breast cancer. Rather, it seems that the Wilkerson subtypes are driven by a complex interplay of multiple, heterogeneous biological processes. Under these circumstances, we would not expect to find factors that represent the subtypes directly, but that the factors should capture the underlying biological processes.

**Fig 6.**
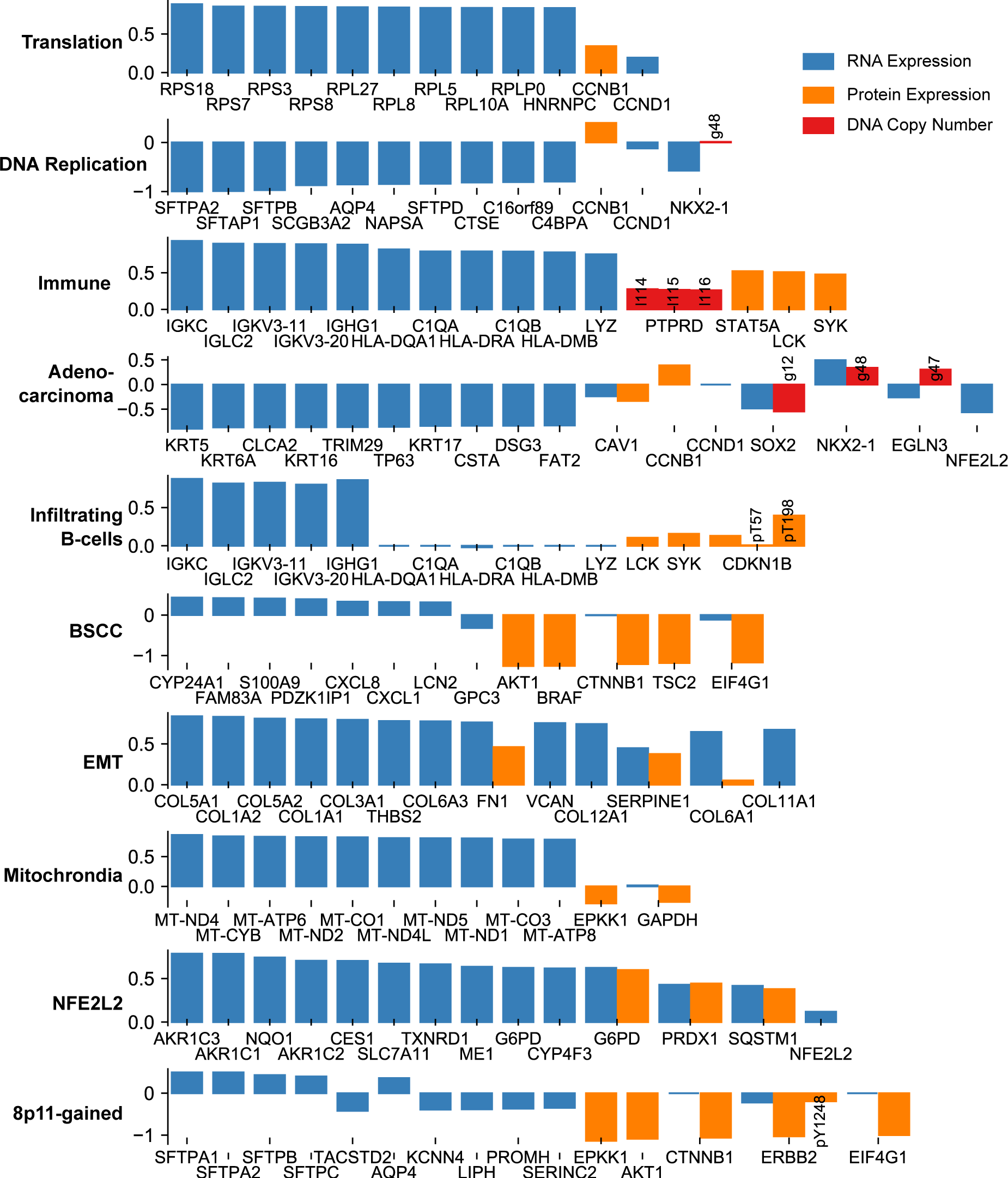
The strongest SFA coefficients for the lung cancer data set for each of the three data types and all ten factors. Height of the bars shows the values of the coefficients. If a gene is shown we show all coefficients of that gene in the model. RNA expression coefficients are shown in blue. Protein expression coefficients are shown in orange. Any modifications of an epitope are noted in a short text description. pX = phosphorylated at residue X. DNA copy number coefficients are shown in red. Numbers refer to the recurrently aberrated loci in S3 Table. Recurrent gains are prefixed with a g, losses with an l.

**Fig 7.**
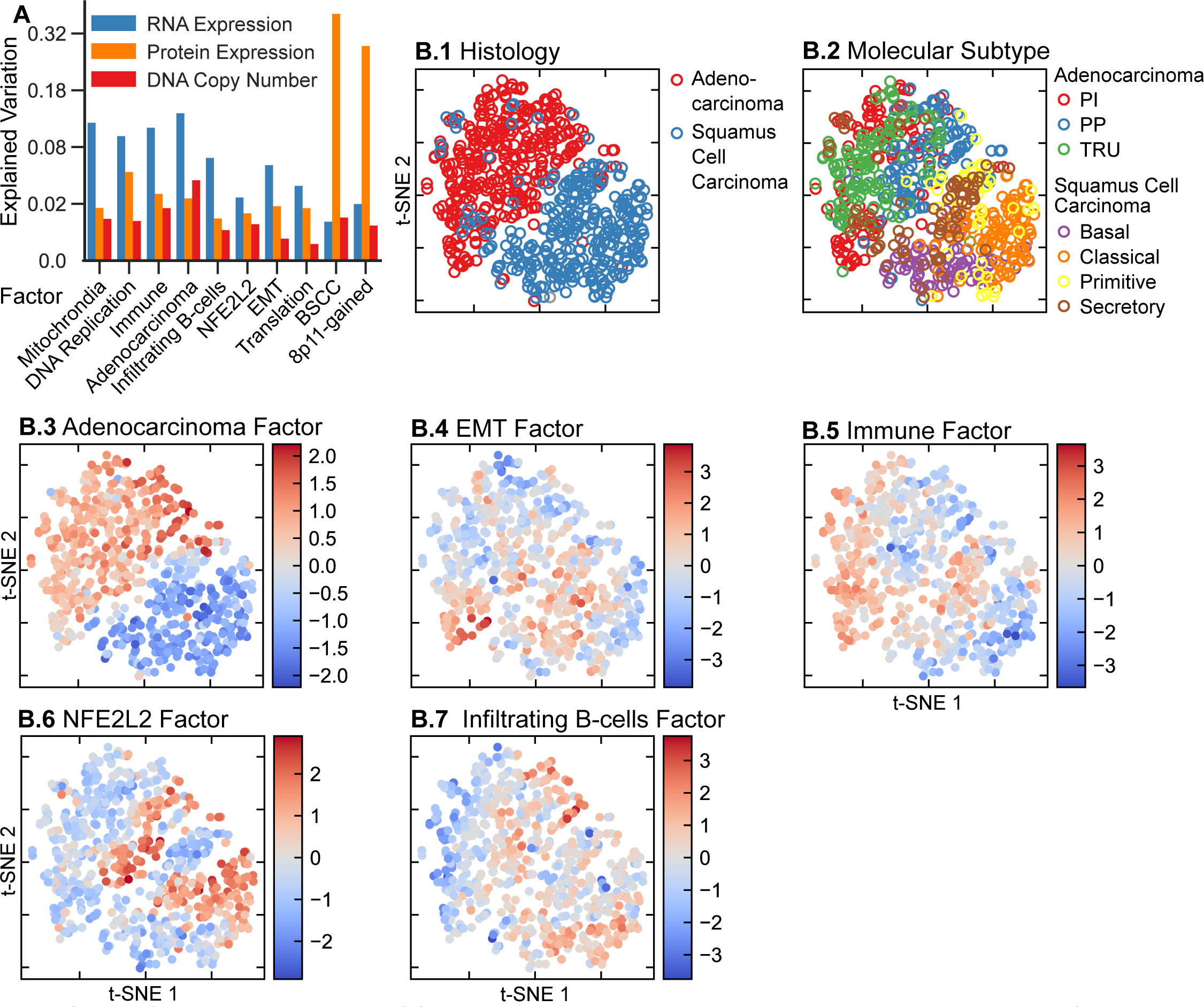
Sparse-factor analysis on the TCGA lung cancer dataset. A: Explained variance per data type and factor. B: B.1 shows the t-SNE map of all lung tumors with red denoting the Adenocarcinomas and blue the Squamous Cell Carcinomas. With the tumors in the same positions as in B.1, B.2 depicts the subtyping as proposed by Wilkerson and colleagues [21, 22]. The remaining panels show the tumors in the same positions as the first two maps, but colored according to the value of the represented factor in each tumor.

**Fig 8.**
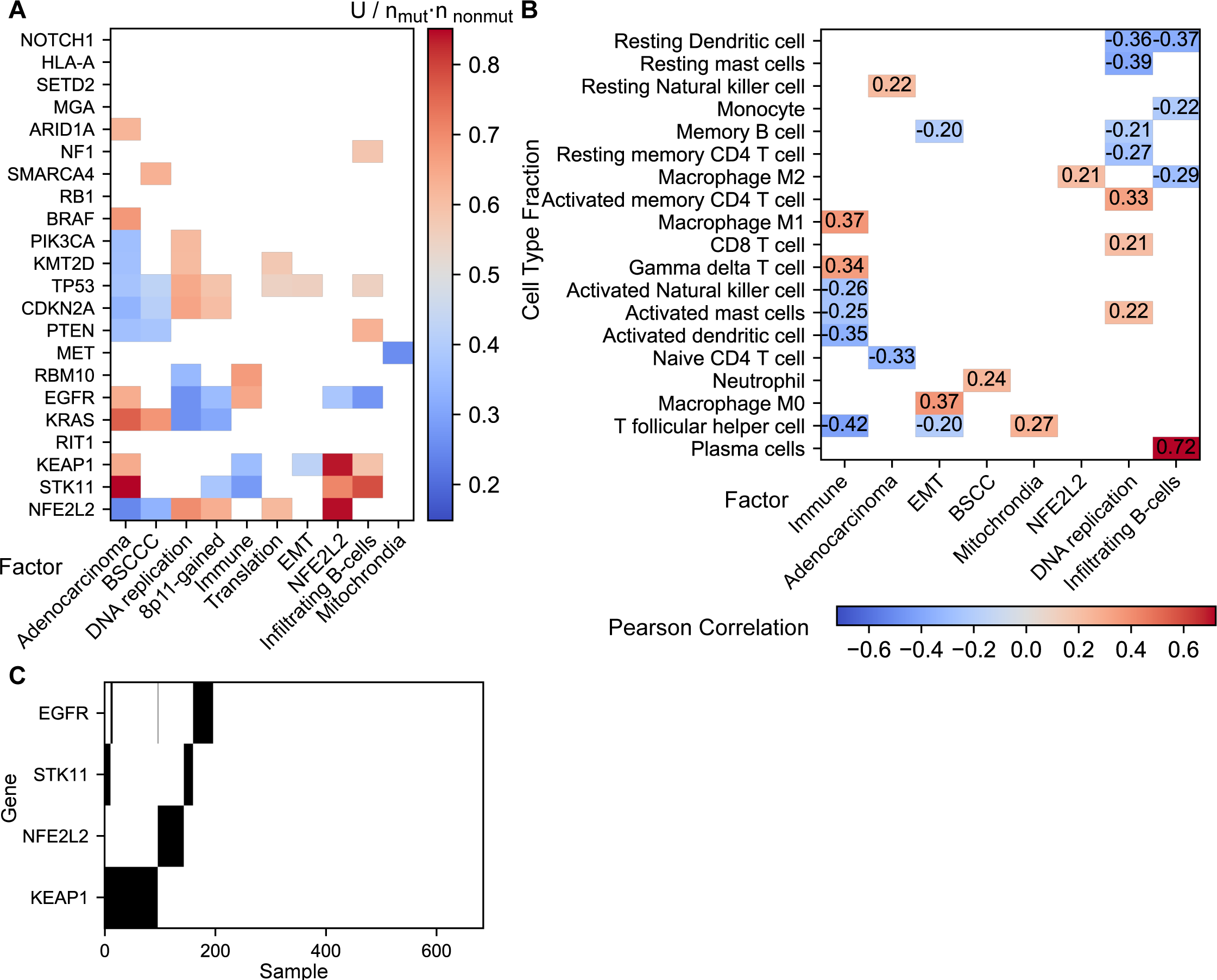
Mutations and immune infiltration in lung cancer. A: Mann-Whithney U statistic of the factor values between tumors with and without a mutation in a gene divided by the product of the number of tumors in each group. Only significant (p<0.05) values are shown. B: Pearson correlation (ρ) between the factors and cell type fractions. Only significant correlations (p<0.05, |ρ|>0.2) are shown. C: Mutations in genes of the NFE2L2 pathway and STK11 (black: tumor is mutated in this gene).

**Fig 9.**
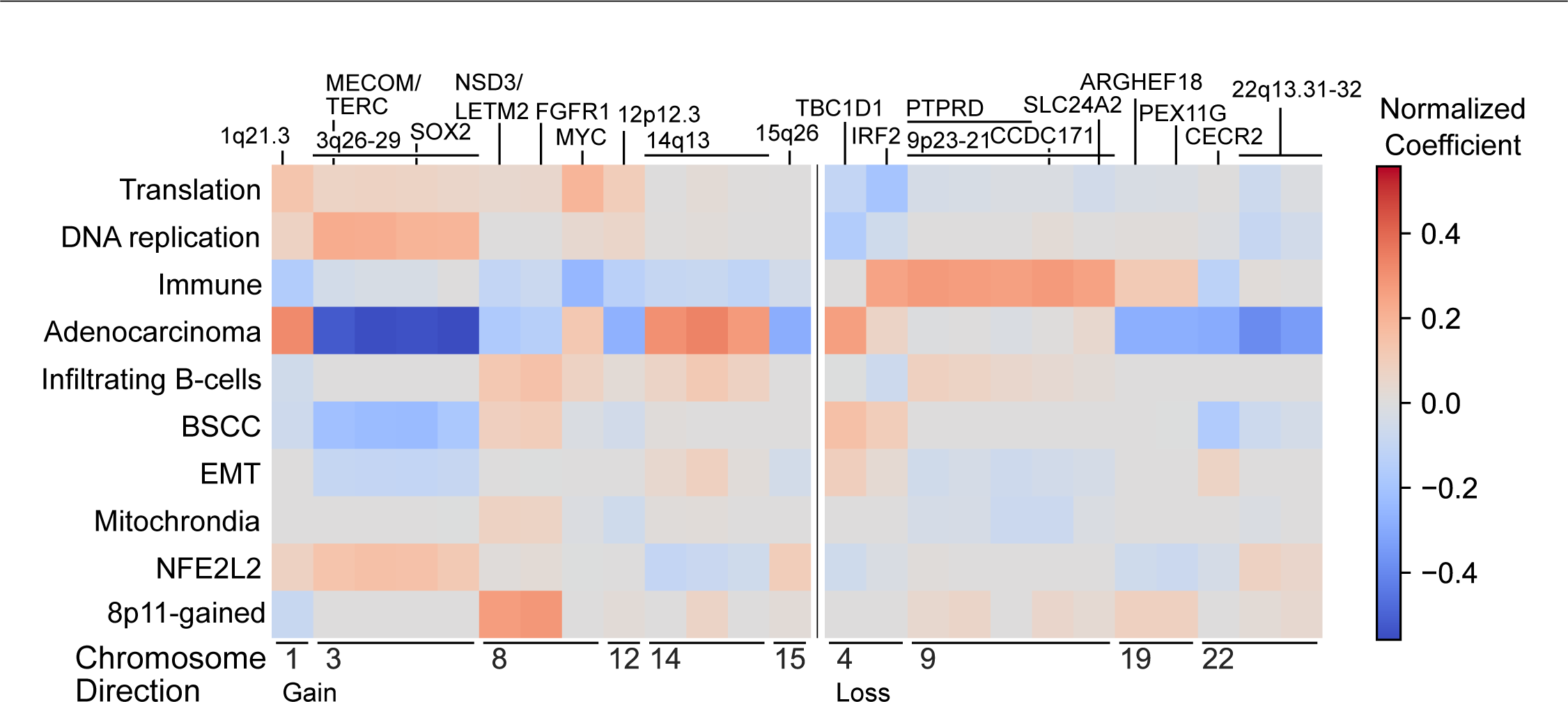
Copy number and factors in the TCGA lung cancer dataset. Normalized coefficients representing the contribution of DNA copy number abberations to the factors. Specifically, the coefficients represent the contribution of recurrently gained (left) or lost (right) copy number regions identified by RUBIC to the factors represented in the rows. Recurrently aberrated copy number regions are annotated with chromosomal bands or putative driver genes in the region.

Adenocarinoma and squamous cell carcinoma were subtyped separately, so we have a set of subtypes for each. The adenocarcinoma subtypes are termed 'terminal respiratory unit’ (TRU), ‘proximal-inflammatory’ (PI) and ‘proximal-proliferative’ (PP). The TRU subtype is characterized by highly expressed asthma, excretion and surfactant genes, the PP subtype with the overexpression of defense response genes (chemokines) and the PI subtype with high expression of DNA-repair genes [22]. The Wilkerson squamous cell carcinoma subtypes are termed ‘Basal’, ‘Classical’, ‘Secretory’ and ‘Primitive’. The Basal subtype has been associated with cell adhesion and epidermal development. Wilkerson and colleagues reported that the classical subtype is related to xenobiotic detoxification, that the Secretory subtype shows association with the expression of immune related genes, NKX2-1 (TTF1), MUC1 and surfactant genes. It is interesting to note that although the secretory subtype is a squamous cell carcinoma subtype, it shows intermediate scores in the Adenocarcinoma factor (Fig 7B.2 and B.3, and S10 Fig), which is consistent with the fact that NKX2-1 is expressed in both the Secretory subtype and in adenocarcinomas. Finally, the Primitive subtype has previously been associated with proliferation and DNA processing and repair.

### Adenocarcinoma and squamous cell carcinoma

The differences between adenocarcinoma and squamous cell carcinoma are captured by the Adenocarcinoma factor, which is high in adenocarcinomas and low in squamous cell carcinomas. Interestingly, in the primitive subtype of squamous cell carcinoma, tumors with high and low values for this factor exist (Fig 7B.2 and B.3, and S10 Fig). In addition, copy number is dominated by a particularly strong negative association with the gain of Chromosome 3q26-29, implying that the adenocarcinomas show absence of gains while squamous cell carcinomas carry this gain (Fig 9). We also observe the expected associations between this factor and mutations in NFE2L2, STK11, KEAP1. KRAS, EGFR, PTEN, TP53, PIK3CA, BRAF and ARID1A (Fig 8A). As expected, the Adenocarcinoma factor is high in all the Wilkerson adenocarcinoma subtypes, as compared to the squamous cell carcinoma subtypes. More specifically, the Adenocarcinoma factor is very low in the Basal and Classical subtypes but higher in the Secretory and Primitive subtypes.

### Two factors strongly associated with RPPA: BSCC and 8p11 gained

The largest fraction of the variance explained by the BSCC and 8p11-gained factors is associated with the RPPA data (Fig 7A). In fact, the amount of variation explained in the RPPA data by these two factors is the highest across all factors and all data types. The 8p11-gained factor is associated with increased gain of recurrent aberrations containing the genes NDS3, LETM2 and, FGFR1, which are located on the 8p11 chromosomal band (Fig 9), and shows high scores in the PI subtype. As expected, the BSCC factor shows, in general, higher scores within the Basal subtype as compared to other squamous cell carcinoma subtypes (S7 Fig and S10 Fig). Specifically, the BSCC factor shows intermediately strong association with the basal subtype of squamous cell carcinoma (AUC=0.59).

### DNA replication factor

The DNA replication factor captures two biological processes. Pathway analysis reveals a positive association of this factor with pathways involved in DNA replication, implying that genes annotated to these pathways are enriched in tumors with high levels of this factor. According to the Cibersort analysis, this factor is positively correlated with signatures representing activated memory CD4 T-cells and negatively correlation with signatures representing resting mast and resting dendritic cells (Fig 8B). Conversely, this factor shows negative association with the protein expression of the transcription factor NKX2-1. As expected, NKX2-1 targets (SCGB3A2 and AQP4) are amongst the top-ranking genes in terms of the magnitude of their negative RNA expression coefficients. Other genes with very large negative RNA expression coefficients for this factor include the surfactant proteins (SFTPA1, SFTPA2, SFTPB, SFTPC, SFTPD) which are co-expressed with NKX2-1 in the alveolar cells of the lung [33].

The DNA replication factor has low scores in tumors of the TRU subtype (S7 Fig and S10 Fig), which is consistent with the association of this subtype with high expression of surfactant proteins. Compared to the TRU subtype, the DNA-replication factor shows high scores in the PI subtype, but not at the same level as in the Primitive and Classical squamous cell carcinoma subtypes. Since the Secretory subtype shows association with surfactant proteins, the DNA-replication factor shows, as expected, low scores for the tumors of this subtype (S7 Fig and S10 Fig).

### Immune factors: Immune and Infiltrating B-cells

FuncSFA identifies two immune related factors: Immune and Infiltrating B-cells (IB-cells). Pathway analysis of the Immune factor shows enrichment for interferon associated signatures and T-cell pathways. This association is confirmed by high RNA expression coefficients of immunoglobulins, members of the histocompatibility complex (HLA) and members of the complement system (Fig 6). According to the Cibersort analysis, the Immune factor correlates negatively with the signatures representing T-follicular helper cells and activated dendritic cells while it positively correlates with signatures of γδ T-cells and M1 macrophages (Fig 8B). In addition, this factor is associated with an increased loss of 9p23-21. The Infiltrating B-cells factor shows strong association with interferon associated signatures and T-cell pathways in the gene-set enrichment analysis. However, this association is weaker than for the Immune factor. When considering the RNA expression coefficients of this factor we see large coefficients for immunoglobulins, but in contrast to the Immune factor, neither the HLA nor the complement system is represented. This factor has a strong positive correlation with plasma B-cells (Fig 8B) and is associated with a higher mutation rate of STK11 (Fig 8A).

As the PP subtype shows overexpression of defense response genes (chemokines), we observe, consistent with this finding, that the infiltrating B-cell and Immune factors score highly in this subtype (Fig 7B.5 and Fig 7B.7). In contrast, the Infiltrating B-cell factor scores lowly in the TRU and PI subtypes. The Immune factor shows high scores in in the TRU, Classical and Secretory subtypes. Interestingly, the Basal subtype shows high levels of the Infiltrating B-cell factor, but no distinctive pattern for the Immune factor (S10 Fig).

### Mitochrondial, Translation and EMT factors

We identified a number of factors with a clear functional profile, but where additional research is required to understand their role and relevance in lung cancer. First, the Mitochondrial factor associated with genes encoded on the mitochondrial DNA and GAPDH protein expression (Fig 6). Second, the Translation factor which shows enrichment for translation pathways and signatures of the cell cycle (S4 Fig). This association is also strongly supported by the large coefficients of genes encoding ribosomal proteins (Fig 6). Finally, as in breast cancer, we also identified an EMT factor in lung. As in breast, the EMT lung factor is strongly correlated with the Anastassiou signature across the tumors (ρ=0.73). Regarding the Wilkerson subtypes, the TRU subtype shows low scores for the EMT and Translation factors while the Translation factor is high in the Primitive subtype.

### NFE2L2

The NFE2L2 factor shows association with NFE2L2 targets in the gene-set enrichment analysis, suggesting this factor captures the activation of NFE2L2. This conclusion is confirmed by a strong association of this factor with mutations in NFE2L2 and in its inhibitor KEAP1 (Fig 8A). Interestingly, the factor is depleted for EGFR mutations, although EGFR activates the NFE2L2 pathway by inhibiting KEAP1 [34]. Still, these three genes in the NFE2L2 pathway show a mutually exclusive mutation pattern (Fig 8C, p<0.05, DISCOVER groupwise test [35]), suggesting that they independently activate the NFE2L2 pathway effecting the common downstream gene expression changes captured by this factor. So, this factor could be a readout of NFE2L2 pathway activation, which in some conditions has been suggested to increase resistance to EGFR inhibitors

In addition to the clear NFE2L2 association, this factor also shows large values for the coefficients for genes related to xenobiotic detoxification, such as CES1, CYP4F3 and CYP4F11. Hence, it is not surprising that the NFE2L2 factor shows high scores in the Classical subtype, which is also associated with this process.

In summary, we have identified factors that do show overall association, but not very clear one-to-one correspondence with the previously defined Wilkerson subtypes. Our factors do capture the major lung cancer subtypes (Adenocarcinoma and Squamous cell carcinoma), and identified processes that occur in both of these types. These include epithelial to mesenchymal transition, two immune processes, and, most interestingly, a factor that reports the activity of the NFE2L2 pathway. The latter may have interesting therapeutic consequences.

## Discussion

In both breast and lung cancer the sparse-factor analysis was able to recover known biology. Additionally, we find factors that capture biology that traditional clustering methods have not been able to find, such as the Immune, EMT, and infiltrating B-cells factors. Our results illustrate the need for continuous factors as opposed to discrete clusters. For example, the Luminal-Proliferative factor shows that the distinction between Luminal A and B is not discrete. Also the need for multiple factors per tumor is illustrated by our results. For example, the Immune and EMT factors in breast cancer are active across all subtypes. Finally, our results show that a complete molecular characterization requires multiple data types. Although all non-technical factors are represented in the RNA expression, the 8q-gained and Luminal Proliferative factors in breast cancer are mostly copy number driven. Also, the protein expression contributes to key factors such as the ER and HER2 factors.

Interestingly we have found the same cancer cell specific EMT factor in both lung and breast cancer. This does not necessarily mean that tumors with high levels of this factor have a large proportion of mesenchymal tumor cells. More likely, this factor represents the activation of a transcriptional program that may eventually lead to a transition to the mesenchymal phenotype. Therefore, the tumors with high levels of this factor can have a higher propensity to undergo EMT. This can potentially be resolved by applying the method pan-cancer including both epithelial and mesenchymal cancers. As the mesenchymal tumors should get a maximum EMT score, this analysis could give an indication how far along the EMT transition the cells in these tumors really are.

Several factors may potentially have an application in predicting a tumors' response to treatment. The NFE2L2 factor might be indicative of response to dimethyl fumarate (DMF) treatment, which targets the NFE2L2 pathway and can inhibit carcinogenesis [36]. Because the factor measures activation of the transcription factor through its downstream targets, it could be more predictive than only looking at mutations in the transcription factor itself. This hypothesis could be tested on patient derived xenograft models or cell lines. The Immune factor in breast cancer and the Immune, DNA replication and B-cell infiltration factors in lung cancer are possibly predictive for response to immune checkpoint inhibitors. If the immune system is already highly active, a clinically relevant response might be easier to achieve. These hypotheses can be tested by obtaining RNA expression data from pre-treatment biopsies of patients treated with immune checkpoint inhibitors, scoring these factors based on these data and correlating that with their response to checkpoint inhibitors.

In the lung cancer data set we found two highly correlated factors. Without the sparsity penalties, the algorithm should find orthogonal factors. Orthogonal factors explain the most variance, but the sparsity penalties prevent full orthogonality. An explanation of the highly correlating factors can be that the optimal sparsity penalty is different per factor, where one biological process involves thousands of genes, and another only a few.

In this study we used only the tumors which have all data types available. With increasing tumors sizes and an increasing number of data types this results in a situation where ever growing parts of the data can not be incorporated in an integrative analysis. The EM framework quite naturally allows for missing data, so this might be an interesting future extension.

In addition to the data types we used here, other molecular data types that could be included are DNA methylation and miRNA expression data. As the mutation data includes driver events it would be desirable to include it as well. This would require transforming the mutation data such that it has approximately Gaussian error. One way to do this is by smoothing mutations over an interaction network [37]. The inclusion of higher level tumor phenotypes, such as features obtained from MR imaging or pathology slides, could guide the method towards finding biological processes that lead to clinically relevant differences.

We have shown FuncSFA is able to find biological processes that are active across otherwise very different tumors, such as the ER+ and ER− subtypes in breast cancer. Applying this method to a pan-cancer cohort might find biological processes that are activated in a large number of tumor types and provide insight into how tumors of different origin relate to each other.

In summary, we have shown that FuncSFA is able to integrate at least three data types. It identifies continuous-valued factors that could be simultaneously active in the same tumor removing the necessity to assign tumors to clusters. Our results illustrate the advantage of factors over clusters with several biologically or clinically relevant examples. The identified factors represent the heterogeneity both within and between cancer types, and represent the activation of biological processes in a patient specific manner. Considering the more complete and fine-grained characterization they allow, these factors could benefit the personalized treatment of cancer.

## Methods

The FuncSFA method consists of two steps. In the first step, a sparse-factor analysis integrates multiple data types into a small number of factors. In the second step, the factors are linked to existing knowledge of biology by doing a gene-set enrichment analysis.

### Sparse-Factor Analysis Model

The data of a single data type *i* with *N* tumors and *n_i_* features is represented in a *n_i_* × *N* data matrix *X_i_*. The data matrices of *t* data types are stacked together into a *n* × *N* data matrix 
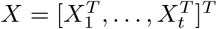
. This data matrix is factorized into a *n* × *k* coefficient matrix *B* and *k* × *N* factor matrix *Z*, where the number of factors *k* is much smaller than the total number of features *n*.

The probabilistic model has been described previously by Shen and colleagues. [8]. We briefly summarize this probabilistic model here. A tumor sample vector **x** with *n* features is explained by factors **z** and coefficients *B* with Gaussian error

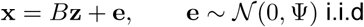

where the residual variance Ψ is zero off the diagonal and equal for features of the same data type. When the factors are taken to be normally distributed

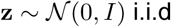

the complete data log-likelihood is:

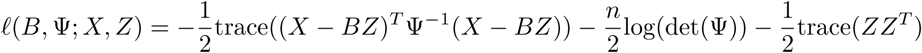

where *X* = [**x**_1_, …, **x***_N_*] and *Z* = [**z**_1_, …, **z***_N_*]. The coefficients are made sparse by a elastic net penalty:

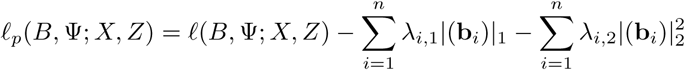

where 
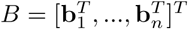
. The penalties *λ*_1,*i*_ and *λ*_2,*i*_ are kept the same for all features in a data type.

### Sparse-Factor Analysis Algorithm

To optimize the penalized complete data log-likelihood we use an iterative algorithm, improving over the iCluster2 algorithm [11] at several key points. It consists of the following steps.

1. Initialize the estimated coefficients *B*̂ and residual variance Ψ̂ by the loadings of a principal component analysis (PCA).
2. Calculate the expectation *E*[*Z|X*] and covariance *E*[*ZZ^T^*|*X*] of the factors given the current *B*ˆ and Ψ̂. In contrast to iCluster2 *E*[*Z|X*] and *E*[*ZZ^T^ X*] are then scaled to have unit variance per factor. This prevents the regularization penalty from forcing large factor values to compensate for small coefficients.
3. Estimation of coefficients is done differently from iCluster2. We use the coordinate descent scheme for solving elastic net as used in glmnet [15]. A coefficient of a single factor *i* and a single feature *j* is updated given the current *E*[*Z*|*X*], *B*̂ and Ψ̂:

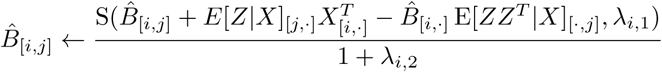

where S is the soft-thresholding operator S(*b*, *l*_1_) = sign(*b*)(|*b*| − *l*_1_)_+_. In a single iteration of the main algorithm the coefficients are updated once in sequential order.
4. Residual variance is estimated per data type:

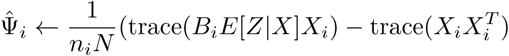

for data type *i*.
5. Repeat from step 2 until convergence, with a maximum of 5.000 iterations. We consider the algorithm converged when the average absolute change in reconstruction error over the last 10 iterations is smaller than 10^-6^.

### Parameter Selection

To find the optimal penalty weight parameters *λ*_*i*,1_, *λ*_*i*,2_ for the L1 and L2 penalties respectively, we reparameterize.

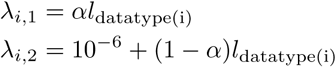

where `datatype(i)` gives the data type of feature *i*. We then do a grid search for the *l* penalties giving the highest Bayesian information criterion (BIC), keeping *α* fixed at 0.9. The effective number of parameters of the penalized coefficients, used for calculating the BIC, is calculated using the method of Tibshirani and Taylor [38].

### Pathway Analysis

We employ a gene-set enrichment analysis tailored to the results of sparse-factor analysis. For this we use gene-set enrichment analysis [10] with a modified gene ranking method. Gene ranking is done as follows. First the factors *Z* are regressed on the RNA expression data containing all genes. Second we get correlation-like coefficients by dividing the regression coefficients of a gene by the standard deviation of that gene. The genes are then ranked per factor based on their correlation-like coefficient.

### Preprocessing

RNA expression data from RNA sequencing was preprocessed using limma-voom [39]. We selected the 1000 genes with the highest median absolute deviation. The mean variance-trend in RNA sequencing data was taken into account by multiplying the RNA expression with the precision weights obtained from voom. DNA copy number was sampled at regions with recurrent copy number aberrations. Recurrently aberrated copy number regions were taken from the RUBIC paper [18] for breast cancer or obtained by running RUBIC for lung cancer. To obtain the copy number of region in a tumor, we took the median copy number of all segments in the tumor sample overlapping that region. This results in a sample by region matrix with copy number data. Protein expression data from RPPA was used directly as processed by TCGA. To remove differences in variance between data types we divided every data type by its standard deviation.

### Additional methods

t-SNE [40] was used to summarize the factors in two non-linear dimensions, yielding a high-level map the factors and other tumor variables can be projected on. These t-SNE maps were calculated using scikit-learn [41].

The explained variance of a factor is calculated by subtracting the summed square error of the model with all factors from the model excluding the factor dividing by the total sum of squares of the data:

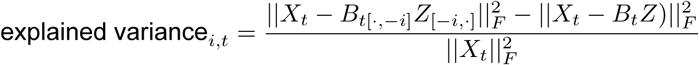

for factor *i* and data type *t* where *B*_*t*[·,−*i*]_ and *Z*_[−*i*,·]_ are the coefficients and factors excluding the factor *i*. The squared Frobenius norm 
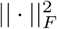
 is the sum of squares of the elements of a matrix.

## Software and data availability

The software for the sparse-factor analysis is available from https://github.com/NKI-CCB/funcsfa. The software for the pathway analysis is available from https://github.com/NKI-CCB/ggsea. The results in this paper are solely based on publicly available data. Breast cancer data was obtained from the TCGA data portal https://tcga-data.nci.nih.gov/docs/publications/tcga/. Lung cancer data was obtained from the Genomic Data Commons Data Portal https://portal.gdc.cancer.gov/.

## Supporting information

**S1 Table. Gene set enrichment of all 10 factors in breast cancer.**

**S2 Table. Gene set enrichment of all 10 factors in lung cancer.**

**S3 Table. Recurrently aberrated loci by RUBIC**

**S1 Fig. Convergence of iCluster, iCluster2 and sparse-factor analysis.**

**S2 Fig. Correlation between the factors of the best solution with a number of factors and the best solution with one factor more.**

**S3 Fig. Histograms of factor values.**

**S4 Fig. Heatmap of GSEA Normalized Enrichment Statistic (Breast)**

**S5 Fig. Heatmap of GSEA Normalized Enrichment Statistic (Lung)**

**S6 Fig. t-SNE maps of breast cancer**

**S7 Fig. t-SNE maps of lung cancer**

**S8 Fig. Scatterplot of coefficients and values of RPPA technical factors in lung.**

**S9 Fig. Boxplots of factors in breast cancer over the PAM50 subtypes.**

**S10 Fig. Boxplots of factors in lung cancer over the Wilkerson subtypes.**

## Acknowledgments

This research is part of the STW Perspectief programme Population Imaging Genetics (ImaGene) and supported by the Dutch Technology Foundation STW, which is part of the Netherlands Organisation for Scientific Research (NWO), and partly funded by Ministry of Economic Affairs.

